# Truncated Complement Factor H Y402 Gene Therapy Cures C3 Glomerulonephritis

**DOI:** 10.1101/2024.09.17.613471

**Authors:** Lindsey A. Chew, Daniel Grigsby, C. Garren Hester, Joshua Amason, W. Kyle McPherson, Edward J. Flynn, Meike Visel, John G. Flannery, Catherine Bowes Rickman

## Abstract

Patients with both age-related macular degeneration (AMD) and C3 glomerulonephritis (C3G) are challenged by the absence of effective therapies to reverse and eliminate their disease burden. Capitalizing on complement dysregulation as both a significant risk factor for AMD and the known pathophysiology of C3G, we investigated the potential for adeno-associated virus (AAV) delivery of complement factor H (CFH) to rescue C3G in a *Cfh-/-* mouse model of C3G. While past efforts to treat C3G using exogenous human CFH resulted in limited success before immune rejection led to a foreign protein response, our findings demonstrate the capacity for long-term AAV-mediated delivery of truncated CFH (tCFH) to restore inhibition of the alternative pathway of complement and ultimately reverse C3G without immune rejection. Comparing results from the administration of several tCFH vectors also revealed significant differences in their relative efficiency and efficacy. These discoveries pave the way for subsequent development of AAV-mediated tCFH replacement therapy for patients with C3G, while simultaneously demonstrating proof of concept for a parallel AAV-mediated tCFH gene augmentation therapy for patients with AMD.

## Introduction

Complement dysregulation is implicated in both age-related macular degeneration (AMD) and C3 glomerulonephritis (C3G). Given the structural and functional similarities^1,2^ between the eyes and kidney, pathobiology and pathophysiology shared by both organs may be less surprising. AMD is a global leading cause of irreversible blindness^3–5^ affecting millions of individuals through loss of their high-acuity, central vision. Simultaneously, half of all patients affected by C3G develop end-stage renal failure^6^ within a decade of presentation. Patients with C3G are commonly diagnosed with mutations^7–11^ in their complement factor H (*CFH*) gene as a primary cause of their kidney disease. These patients also frequently present with coincident ocular subretinal deposits that are indistinguishable^12–14^ in their composition from characteristic AMD extracellular deposits known as drusen^8,15,16^. Notably, the increased presence of drusen^17–19^ is considered a hallmark of “dry”, non-neovascular AMD, and significant AMD risk is conferred by the CFH genetic variant^20–24^ in which tyrosine is substituted for histidine in amino acid 402 (Y402H). This evidence supports certain shared etiology underlying AMD and C3G, while simultaneously highlighting the potential for treatments that rescue both AMD and C3G disease phenotypes. CFH variants and mutations linked to AMD^24^ and C3G, respectively, motivated our consideration of adeno-associated virus (AAV)-mediated gene therapy as a strategy for treatment of these diseases.

Delivery of AAV-mediated gene therapy to the outer retina remains challenging. Current subretinal injection methods in mouse models are limited by the iatrogenic damage induced by the procedure’s requirement for perforation of the neural retina. This obstacle eliminates the opportunity for meaningful assessment of visual function in mice following treatment, thereby hindering the development and testing of gene therapy treatments for AMD. However, systemic delivery of AAV-mediated gene therapy by tail vein injection does not face the same challenge. This provides an ideal framework for testing the therapeutic efficacy of an AAV-mediated gene therapy strategy through an alternative model of disease by treating C3G and eliminating renal disease.

We previously published the design of several AAV gene therapy vectors to support human CFH Y402 replacement in *Cfh* knock-out (*Cfh-/-*) mice^25,26^, which not only provides a straightforward platform for measuring successful gene replacement but also serves as a well-established mouse model of C3G^26^. CFH consists of twenty short consensus repeats (SCRs), with several SCRs known for their roles in C3- or glycosaminoglycan (GAG)-binding^27,28^ (**Figure 1**). We created truncated versions of human CFH (tCFH) named for the total number of SCRs expressed in each truncated CFH cDNA. We initially compared the efficiency of AAVs^25^ inducing expression of full-length CFH, 18tCFH, and factor H like 1 (FHL-1, the endogenous human splice variant of CFH composed of SCRs 1-7) in *Cfh-/-* mice (**Figure 1**). In untreated *Cfh-/-* mice, CFH deficiency results in complement dysregulation, aberrant C3 tick-over^26,29,30^, and uninhibited consumption of C3 and Factor B (FB) by the formation of the C3 convertase, C3bBb, of the alternative pathway of complement (APC)^31^. We showed that AAV-mediated CFH Y402 replacement therapy in *Cfh-/-* mice restores complement regulation through the APC, leading to detectable recovery of FB levels^25^. Our findings underscored the importance of local synthesis for ocular complement regulation, while simultaneously revealing differences in the efficacy of CFH replacement according to the SCRs expressed following treatment^25^.

**Figure 1.**
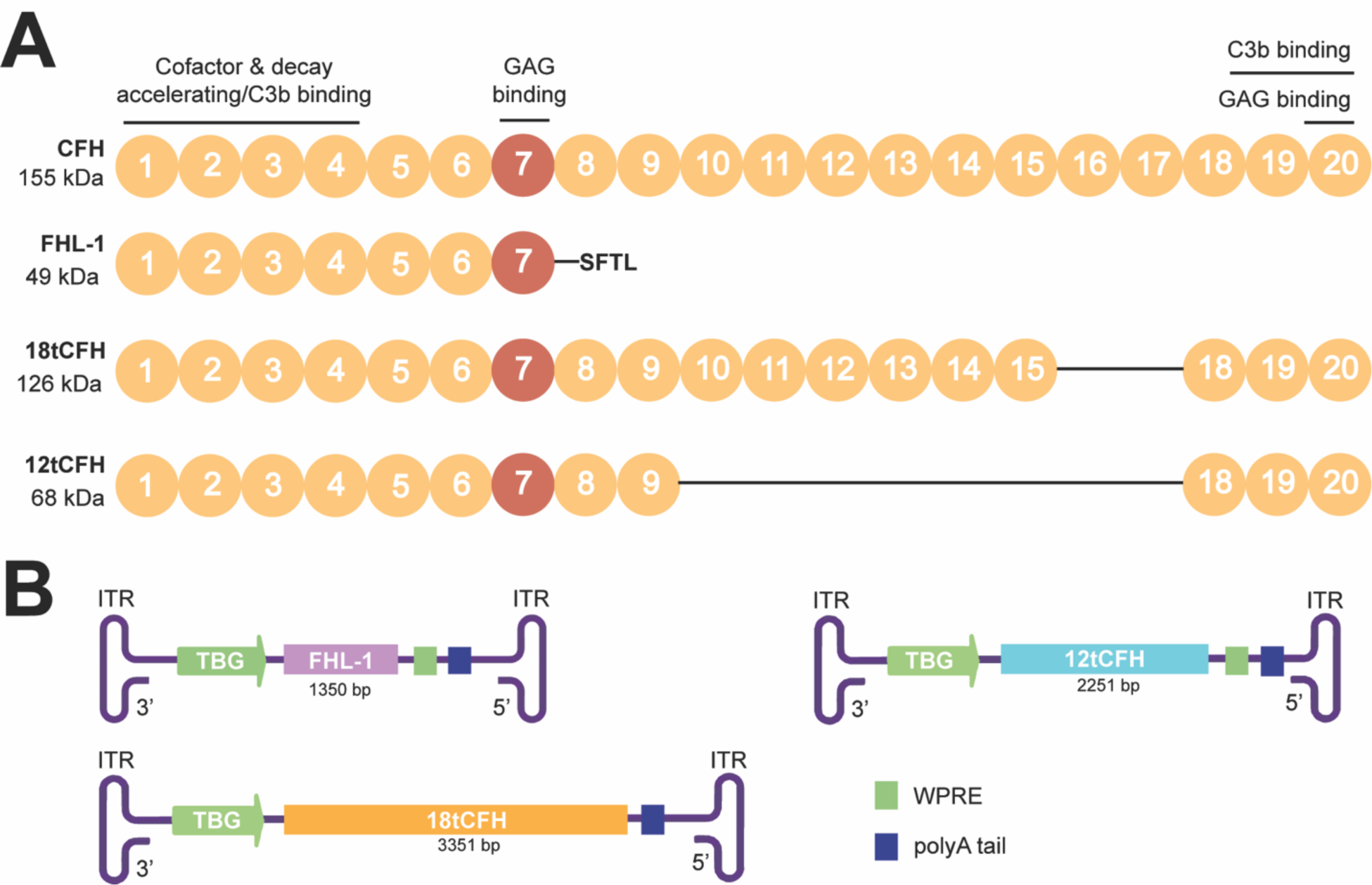
Schematic of CFH and truncated CFH (tCFH) proteins packaged into AAV targeting hepatocytes. (**A**) Full-length CFH (CFH) consists of 20 short consensus repeats (SCRs) including the SCR spanning the AMD-risk variant CFH Y402H AA (red). Key binding domains and known roles are highlighted. The size (kDa) and retained SCRs for each tCFH protein is also shown. Line connecting SCRs denote missing SCRs compared to CFH. GAG, glycosaminoglycan (**B**) Each construct from inverted terminal repeat (ITR) to ITR, with its corresponding promoter (green arrow), *tCFH* cDNA (its size). TBG, thyroxine-binding globin; WPRE, Woodchuck Hepatitis B Post-Translational Response Element and poly A tail.

In this study, we tested the hypothesis that AAV-mediated human CFH Y402 replacement therapy can be used as a treatment to cure C3G in the *Cfh-/-* mouse model. Previous efforts to treat *Cfh-/-* mice by systemic administration of purified human CFH (hCFH) were plagued by long-term immune rejection of the hCFH^32^. In contrast, *CFH-*humanized *Cfh-/-* mice expressing either the human CFH transgenes for the AMD-protective Y402 variant (*CFH- Y/0*) or AMD-risk H402 variant (*CFH-H/H*) of hCFH do not develop C3G^29,33^. Expressing hCFH from birth, these mice also do not mount a foreign protein response to the hCFH^33^. These results show that endogenous hCFH can effectively replace mouse cfh in *Cfh-/-* mice to reverse the C3G phenotype and treat the nephropathy^2,3^. Building on past conceptual achievements in the development of CFH gene therapy^25,32^, this supports the hypothesis that AAV-mediated gene replacement will not induce a foreign protein response like that detected following treatment with exogenous human CFH protein^32^.

Importantly, C3G requires diagnosis by histopathology; more specifically, C3G diagnosis relies on renal biopsy demonstrating evidence of glomerulonephritis^34,35^ with sole or dominant glomerular immunofluorescence staining for C3 of at least two orders of magnitude greater intensity than for any other immune reactant. We relied on this gold standard for C3G diagnosis along with histopathological assessment of the kidney glomeruli^11,35^ in treated *Cfh-/-* mice to verify successful CFH replacement therapy compared to untreated controls. Our findings provide *in vivo* preclinical evidence for AAV-mediated CFH Y402 replacement therapy as a viable treatment strategy to treat and resolve C3G.

## Results

### Systemic, AAV-mediated expression of hepatic tCFH in *Cfh-/-* mice restores intact C3 and reduces C3b through restoration of the APC in the fluid phase

Complete CFH deficiency in *Cfh-/-* mice leads to uninhibited consumption of C3^26^ and cleavage into C3b to support formation of the C3 convertase complex; we hypothesized that AAV-mediated CFH replacement would reduce C3b in the fluid phase by restoring the APC to conserve intact C3. To test this, we designed recombinant AAV constructs optimized for liver expression according to efficiency of the selected AAV serotype and a liver-specific promoter. This design strategy aligned with natural systems where the bulk of systemic CFH is produced by the liver^36^. Specifically, we used AAV8 to efficiently transduce hepatocytes^37^ and a thyroxine-binding globin (TBG) promoter^38,39^ to drive hepatocyte-specific expression of the tCFH cDNAs. Three tCFH AAVs coding for three truncated versions of CFH were used (**Figure 1**). In addition to our previously characterized 18tCFH and FHL-1 AAV expression cassettes, we also designed a 12tCFH AAV construct with a strong likelihood of overcoming the limited packaging capacity of AAVs. Containing 6 fewer SCRs than its 18tCFH counterpart, the 12tCFH AAV construct would be expected to express higher levels of its significantly shorter tCFH gene product^40^. This expectation mirrors observed expression of the FHL-1 construct, which contained 11 fewer SCRs and was expressed at higher levels than the 18tCFH construct^25^ (**Figure 1**). **Figure 1A** highlights motifs from full-length CFH that are expressed in the tCFH proteins, including varying relative SCRs with different known functions. **Figure 1B** depicts the tCFH expression cassettes inserted in AAV8 capsids for use in these studies.

The three AAV8-TBG-tCFH AAVs (**Figure 1B**) were systemically administered to *Cfh-/-* mice of various ages via tail vein injections. Three trials of this experiment were conducted using the viral titer for each iteration of the experiment as shown in **Table 1**. Blood was collected^29,34,35^ 8 to 12 weeks after tail vein injection from treated animals (*Cfh-/-*AAV12tCFH; *Cfh-/-*AAVFHL-1; *Cfh-/-*AAV18tCFH), untreated *Cfh-/-* animals (which serve as C3G-positive controls presenting with kidney disease), and untreated *CFH-H/H* animals (which serve as a C3G-negative control without kidney disease). Plasma was isolated from the blood of each animal. The levels of 12tCFH, 18tCFH, and FHL-1 proteins in the plasma of AAV-treated animals was quantified by Western blotting (**Figure 2A**). A comparison of these immunoblots revealed higher levels of circulating 12tCFH and FHL-1 compared to levels of 18tCFH in treated *Cfh-/-* mice (**Figure 2A**). We assessed restoration of the APC by quantifying levels of C3 and C3b in the plasma of these same animals (**Figure 2B**). While uninhibited C3 tick-over in *Cfh-/-* mice leads to complete consumption of C3, leaving only the cleavage product C3b^26,29,30^, successful AAV-mediated CFH replacement therapy and subsequent restoration of the APC leads to detectable recovery of intact C3 (**Figure 2B**). The ratio of C3/C3b in each lane provides a normalized scale for restoration of the APC despite animal-to-animal variability in circulating levels of complement and components of its pathways (**Figure 2C**). Across treated and untreated animals, examining the ratio of C3/C3b in fluid phase (**Figure 2C**) highlights the concentration-dependent recovery of the APC based on the levels of tCFH or FHL-1 (**Figure 2B**). Although low levels of circulating 18tCFH are observed in *Cfh-/-*AAV18tCFH mice, proportional levels of intact C3 are still present (**Figure 2A**).

**Figure 2.**
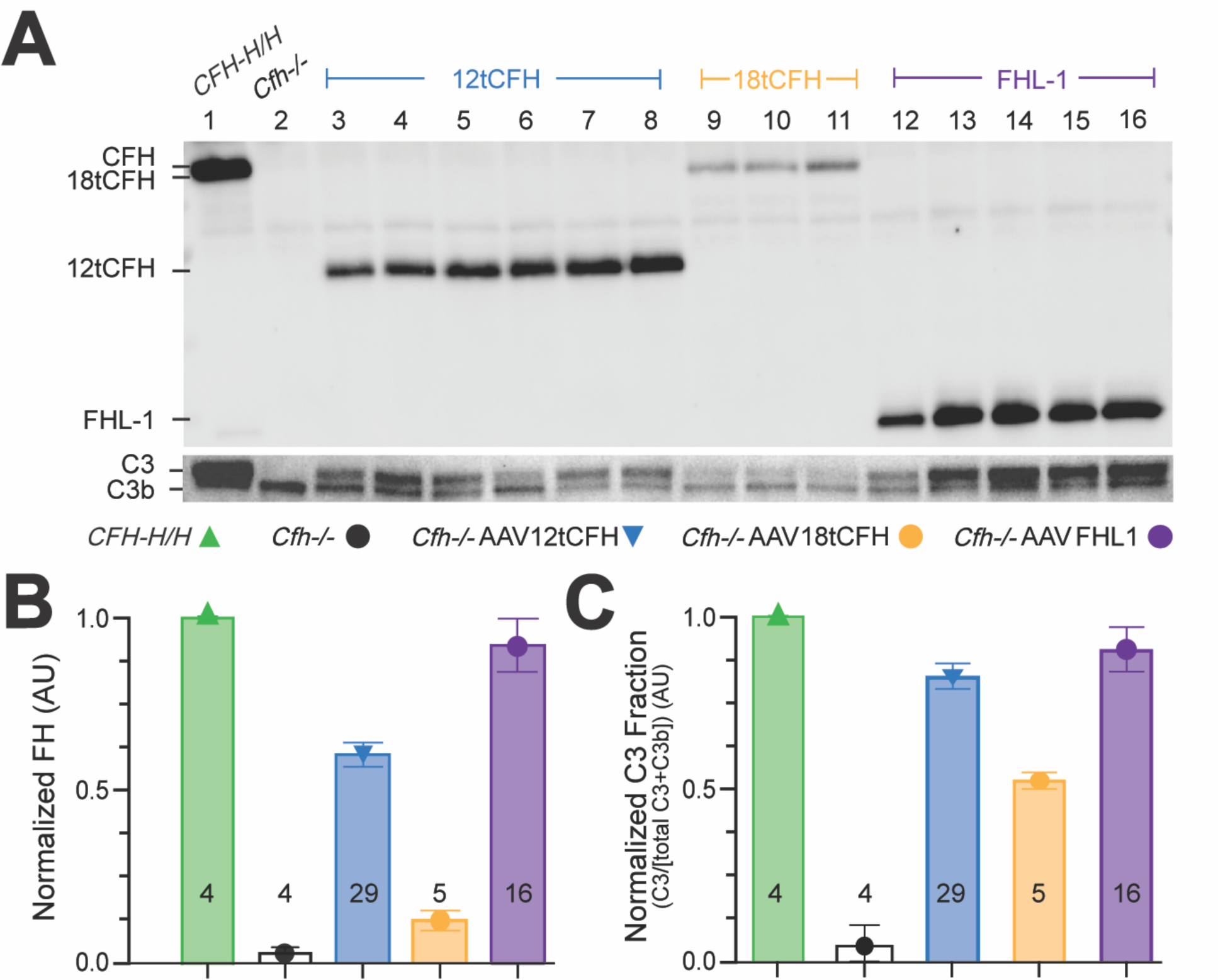
Systemic, AAV-mediated replacement of CFH in *Cfh-/-* mice drives hepatic tCFH expression and reduces fluid phase C3b through restoration of the APC. (**A**) Representative Western blots showing circulating plasma levels of 12tCFH, 18tCFH, and FHL-1 in *Cfh-/-* mice treated by AAV-mediated CFH replacement. (**B**) Bar graph indicating normalized CFH expression in each cohort (± AAV treatment). The number of mice analyzed for each cohort is shown at the base of each bar. (**C**) Bar graph of normalized C3 fraction calculated as the C3 fraction of the total C3 + C3b for each animal and normalized against *CFH-H/H,* C3G-negative control mice. The number of mice analyzed for each cohort is shown at the base of each bar.

**Table 1.**
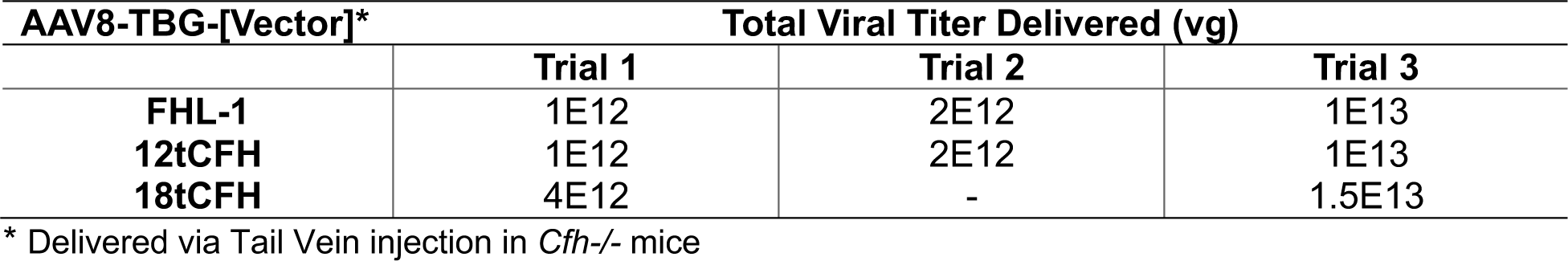
AAV Vectors and Viral Titers.

### AAV-mediated CFH replacement reduces C3 deposit accumulation and resolves C3G

Renal biopsy followed by histology and glomerular immunofluorescence are both necessary and sufficient components of C3G diagnosis^34,35^. Previous efforts have characterized the *Cfh-/-* mouse model of C3G by these measures, while simultaneously demonstrating that uncontrolled C3 activation *in vivo* is essential for the development of C3G membranoproliferative glomerulonephritis^26,29^. This requirement is supported by evidence of normal renal function and histology in double knock-out mice (*Cfh-/-:Bf-/-)* with both CFH-deficiency and factor B-deficiency (*Bf-/-)*, which prevents formation of the C3 convertase C3bBb^26^ through the APC. In *Cfh-/-* mice, uninhibited consumption of C3 leads to excesses of C3b/iC3b in the fluid phase, contributing to the accumulation of significant C3b/iC3b deposits in the glomeruli, with subsequent development of kidney disease and, ultimately, renal failure^32,35,41^. We hypothesized that successful tCFH replacement therapy would significantly reduce the accumulation of C3 deposits in treated *Cfh-/-* mice.

To test this, kidneys were harvested for analysis from treated animals (*i.e. Cfh-/-* AAV12tCFH; *Cfh-/-*AAVFHL-1; *Cfh-/-*AAV18tCFH), untreated positive-control (C3G-positive, C3G+) *Cfh-/-* animals, untreated positive-control *CFH-H/0* animals (transgenic mice that are also C3G+ because they express full-length hCFH H402 at only ∼10% of wild type mouse levels^29^), and untreated negative-control (C3G-negative) *CFH-H/H* animals^33^. After sectioning and immunolabeling with anti-C3, we analyzed glomeruli from these kidneys for C3 immunofluorescence (**Figure 3A**). We conducted a masked analysis of glomeruli from three kidney sections per mouse across 16-24 mice for each *Cfh*-/-AAV treatment condition, and we graded each glomerulus as C3-positive (C3+, with C3 immunolabeling present) or C3-negative (C3-, without C3 immunolabeling present). This analysis allowed us to determine the ratio of C3+ glomeruli as a fraction of the total number of glomeruli in the sections captured (**Figure 3B**). Our results demonstrate that AAV-mediated tCFH replacement with 12tCFH and 18tCFH correspond with significantly lower ratios of C3+ glomeruli/total glomeruli compared to untreated *Cfh-/-* mice (**Figure 3B**). While AAV-mediated FHL-1 replacement therapy also significantly lowered the C3+ glomeruli/total glomeruli ratio, it appeared to be only 50% as effective as 12tCFH or 18tCFH replacement (**Figure 3B**). Further separation of experimental cohorts treated with FHL-1 replacement revealed a dose-dependent effect of FHL-1 replacement that strongly corresponded to the viral titer administered by tail vein injection (**Figure 3Avi,vii; 3C**). The apparent dose-dependent clearance of C3 deposits following treatment by AAV8-TBG-FHL-1 mimics the CFH concentration-dependent effect obtained by the clearance of C3 deposits in glomeruli of *CFH-H/H* mice, but not *CFH-H/0* or *Cfh-/-* mice (**Figure 3A, B**). These findings confirm our previous results^25^, suggesting that 12tCFH and 18tCFH work more efficiently than FHL-1 in plasma (**Figure 3B**).

**Figure 3.**
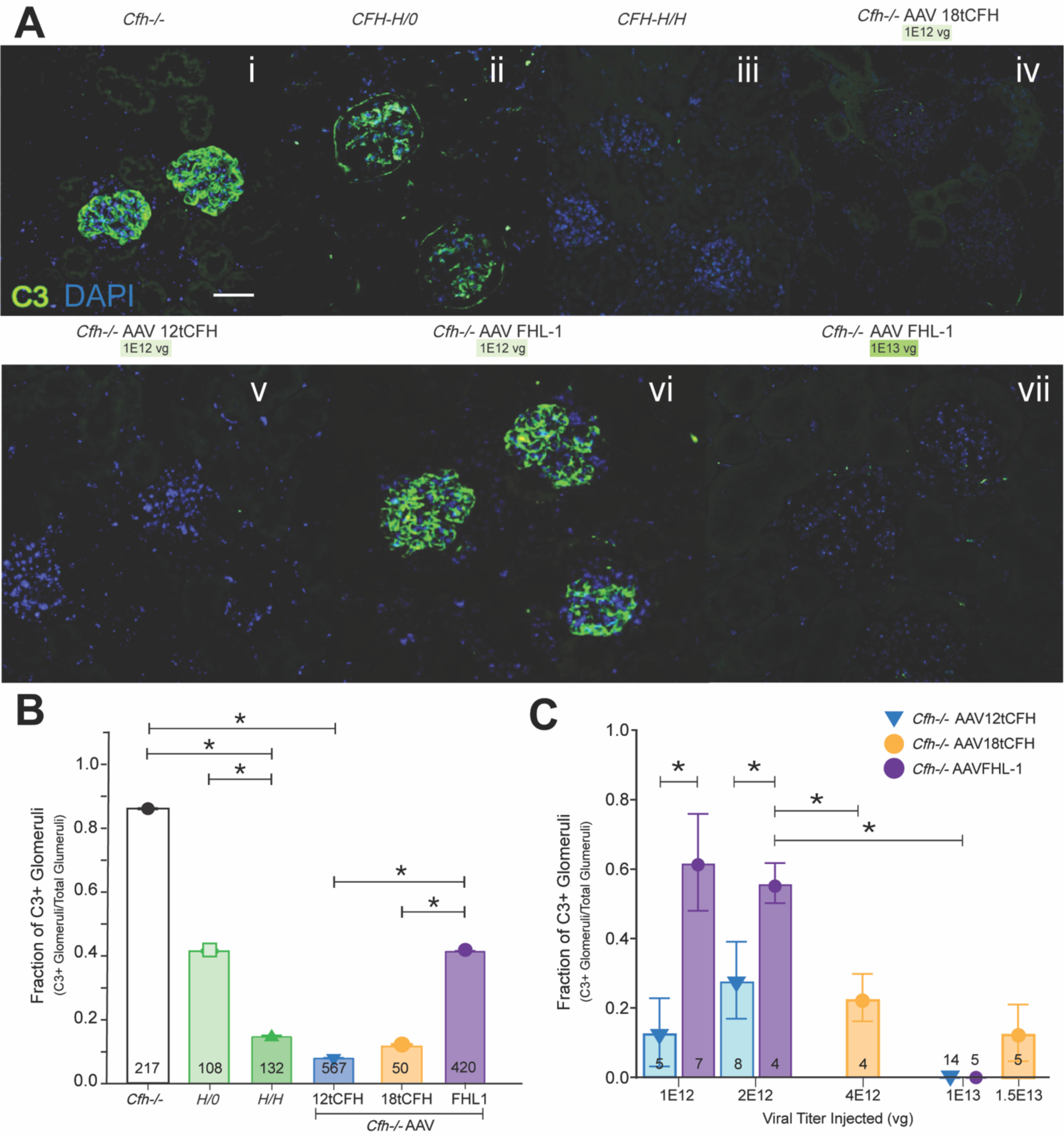
AAV-mediated CFH replacement reduces C3 deposit accumulation and resolves C3G. (**A**) Representative immunofluorescent confocal imaging of glomeruli from kidney sections of mice according to genotype indicated and AAV treatment (viral titer delivered) as shown. Scale bar is 100 microns. (**B**) Bar graph indicating the fraction of C3-positive (C3+) glomeruli out of the total glomeruli for which confocal imaging was obtained. The number of glomeruli analyzed for each cohort is shown at the base of each bar. *Cfh-/-* n=9; *CFH-H/*0 n=6; *CFH-H/H* n=9; *Cfh-/-* AAV12tCFH n=29; *Cfh-/-* AAV18tCFH n=9; and *Cfh-/-*AAVFHL-1 n=16. Unpaired t-test was used for comparison between treatment cohorts, *p<0.01. (**C**) Bar graph analyzing only AAV-treated *Cfh-/-* mice and assessing treatment as a function of the fraction of C3+ glomeruli out of the total glomeruli imaged. The number of mice in each treatment cohort is shown at the base of each bar. Unpaired t-test was used for comparison between treatment cohorts, *p<0.01.

We further analyzed this CFH-concentration dependent effect by comparing glomeruli from *Cfh-/-* AAV mice treated across several different doses of total AAV viral genomes (**Figure 3C**). The fraction of C3+ glomeruli was significantly lower in *Cfh*-/-AAVFHL-1 mice treated with the highest dose (1E13 vg) of AAV8-TBG-FHL-1 than in counterparts treated with lower doses of the same AAV (*i.e.* 1E12 vg or 2E12 vg) (**Figure 3C**). In *Cfh*-/-AAV12tCFH treated mice, there was no significant difference in the fraction of C3+ glomeruli across several cohorts that received different doses. These findings also point to a concentration-dependent effect governing clearance of C3 deposits by FHL-1 where lower doses did not resolve the C3G as well, while 12tCFH-mediated clearance of C3 deposits does not appear constrained by the same concentration-dependent limitation.

### AAV-mediated CFH replacement minimizes hallmark histopathological findings in glomeruli and resolves C3G

In the *Cfh-/-* mouse model of C3G, thickening of the glomerular basement membrane (GBM) and mesangial matrix expansion are hallmark characteristics attributed to uncontrolled C3 activation ^26^. As a result, we expected that tCFH replacement would significantly limit the severity of histopathological findings in the glomeruli of treated *Cfh-/-* mice; specifically, we anticipated that GBM thickness and mesangial matrix expansion would not be significantly different from *CFH-H/H* counterparts without renal disease.

Kidney sections were stained with periodic acid Schiff (PAS), to highlight any glomerular damage in treated animals (*i.e. Cfh-/-*AAV12tCFH; *Cfh-/-*AAVFHL-1; *Cfh-/-* AAV18tCFH), untreated *Cfh-/-* animals, untreated *CFH-H/0* animals, and untreated *CFH-H/H* animals. We graded the histological appearance of each glomerulus as previously described^29^ and assigned a holistic score to each animal, as shown in **Table 2**. Our observations highlight the significant reduction of C3G’s hallmark pathologic features in glomeruli of AAV-treated cohorts. Grading scores of *Cfh*-/-AAV cohorts were significantly lower than scores of untreated counterparts (**Figure 4**). Hallmark pathologic features such as GBM thickening and extracellular matrix expansion are visible in the untreated *Cfh-/-*, *CFH-H/0, and CFH-H/H* mice (**Figure 4i-iii)**. These features appear to be resolved or significantly less severe amongst the *Cfh*-/-AAV treated animals (**Figure 4iv-vi**). Comparing the average histopathology grade of each *Cfh-/-*AAV treatment cohort reinforces the capacity of AAV-mediated human tCFH expression to significantly reduce glomerular damage. Despite differences in the circulating quantities of CFH, 12tCFH, 18tCFH, or FHL-1, mice across all *Cfh*-/-AAV treatment cohorts exhibited significantly less GBM thickening or extracellular matrix expansion in their glomeruli compared to their untreated *Cfh-/-* and *CFH-H/0* counterparts. Altogether, the reversal of hallmark, pathologic signs of C3G through AAV-mediated tCFH replacement therapy emphasizes the viability of this strategy to resolve C3G in the *Cfh-/-* mouse.

**Figure 4.**
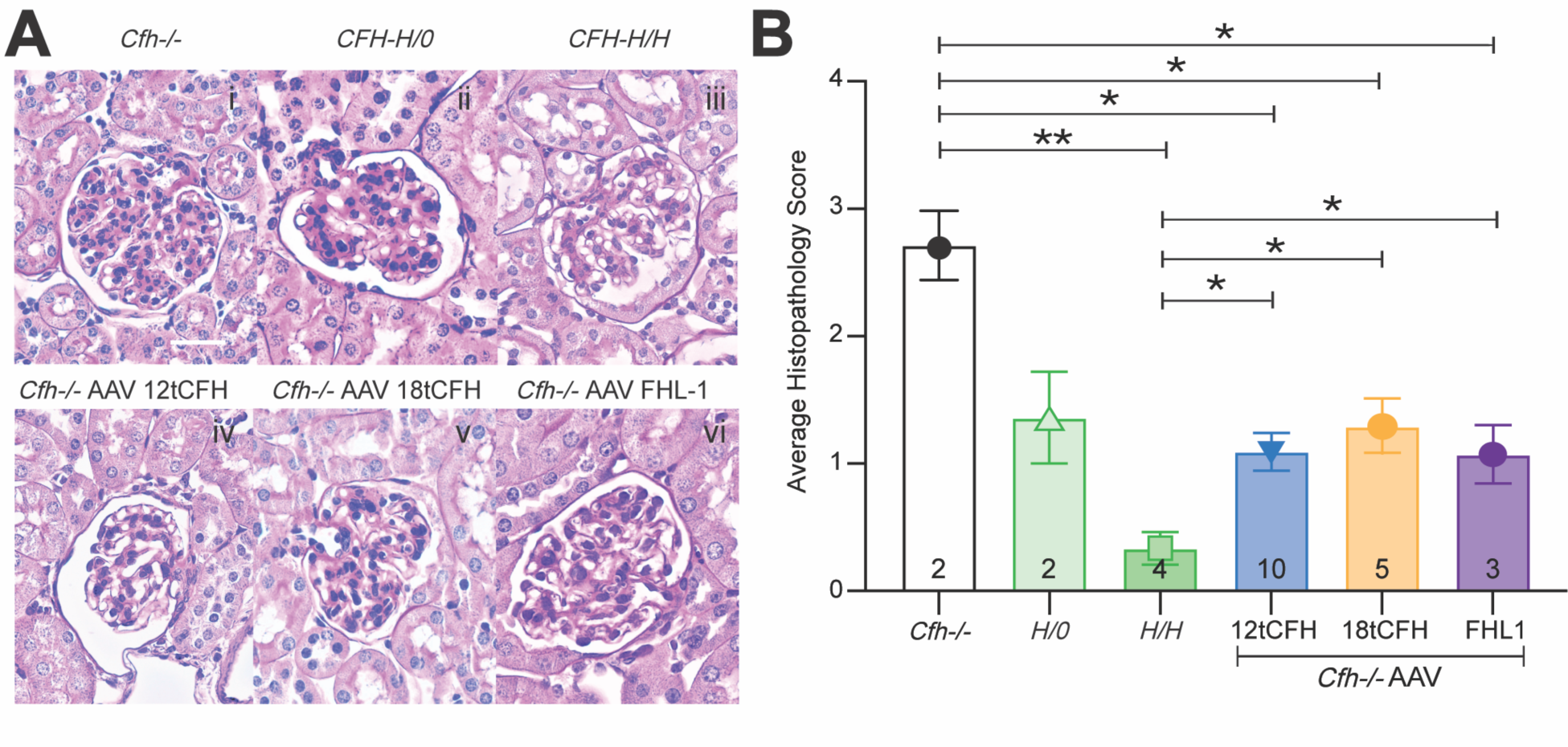
AAV-mediated CFH replacement minimizes hallmark histopathological findings in glomeruli and rescues C3G. (**A**) Representative images following immunohistochemistry using Periodic-acid Schiff (PAS) staining to highlight features of the glomerulus of a kidney. Scale bar corresponds to 20 microns. *Cfh-/-* glomeruli had notable hypercellularity, some mesangial matrix expansion, and basement membrane (BM) thickening. *CFH-H/0* mild hypercellularity, mild BM thickening, mild mesangial matrix expansion. *CFH-H/H* healthy glomeruli. *Cfh-/-*AAV12tCFH (1E13 vg), healthy glomeruli. *Cfh-/-*AAV18tCFH (1.5E13 vg), hypercellularity and mesangial matrix expansion. *Cfh-/-*AAVFHL-1 (1E13 vg), mild hypercellularity and mild mesangial matrix expansion. (**B**) Bar graph comparing masked scores assessing histopathology across mice with and without treatment. The number of mice in each cohort is shown at the base of each bar. Unpaired t-test was used for comparison between treatment cohorts, * p < 0.05, **p<0.01.

**Table 2.**
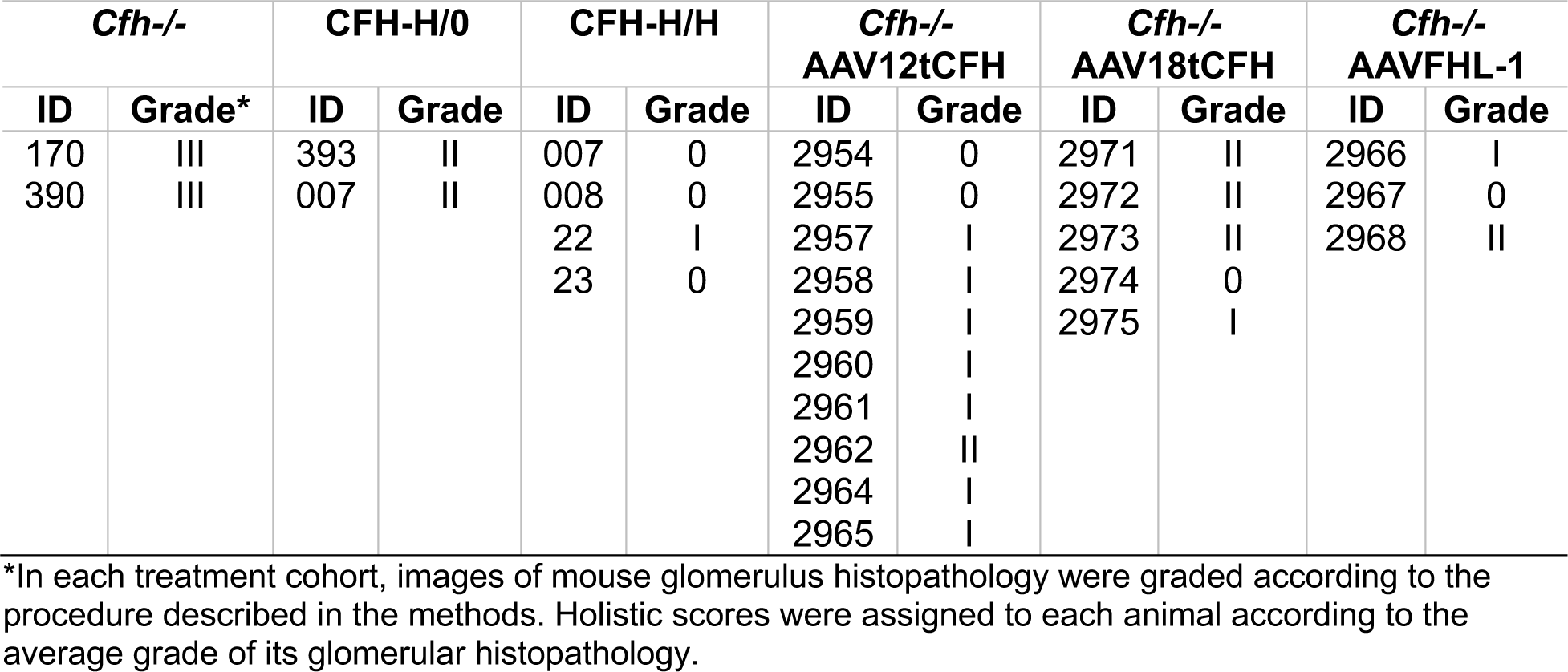
Grading of mouse glomeruli histopathology in each treatment cohort.

## Discussion

C3G develops from uncontrolled C3 activation resulting from CFH deficiency in *Cfh-/-* mice and certain patients^26^. CFH deficiency can arise from loss of function mutations and truncations^42,43^, from inadequate CFH expression^44^, or even from failure of CFH-associated transcription factors^45,46^, and these failures ultimately result in dysregulation of C3 tick-over ^26,29,30^. Although past efforts to restore CFH in the circulation through supplementation with exogenous hCFH^32^ were stymied by activation of anti-human CFH antibodies in the *Cfh-/-* mouse, our findings clearly demonstrate the feasibility of an AAV-mediated gene therapy for CFH replacement to treat C3G. Since the *Cfh-/-* mouse is well-established as a model of C3G^26^, reversing disease in this mouse represents immense promise in the development of a cure for C3G. More specifically, measurable pathologic features of the *Cfh-/-* mouse include glomerular C3 deposit accumulation (**Figure 3**), GBM thickening, and mesangial matrix expansion (**Figure 4**); quantifiably reversing these endpoints following successful treatment strongly suggests progress towards an effective C3G therapy. We show that treatment with each of our AAV vectors (AAV8-TBG-12tCFH, AAV8-TBG-18tCFH, and AAV8-TBG-FHL1) could successfully rescue C3G in this model. Evidence from not only the plasma of treated animals (**Figure 2**), but also the glomeruli of treatment animals underscored the dramatic phenotypic change with treatment (**Figures 3, 4**). Successful treatment through this AAV-mediated strategy also avoided long-term immune rejection and systemic damage, with no evidence of immune infiltration and rejection in the kidneys (**Figure 4**), in contrast to the previously described failure of treatments injecting exogenous hCFH into *Cfh-/-* mice. Specifically, the treatment of C3G in *Cfh-/-* mice, with exogenous hCFH for more than five days, led to glomerular reactivity with anti-C3 antibodies, due to deposition of mouse IgG–hCFH immune complexes^32^. Our twelve-week treatment regimen did not show these staining patterns. The most likely explanation for success and evasion of immune rejection in this strategy includes that an AAV-mediated strategy directing liver-specific expression of tCFH constructs allows for endogenous glycosylation of the gene product. In contrast, exogenous introduction of injected hCFH^32^ includes species-specific glycosylation that is more likely to flag the immune system against the foreign gene product.

However, the AAV treatments were not equal. In the *Cfh-/-*AAV18tCFH cohort, treatment by AAV8-TBG-18tCFH did not induce 18tCFH expression levels as high as 12tCFH and FHL-1 expression achieved by the AAV8-TBG-12tCFH and AAV8-TBG-FHL1 vectors. 18tCFH has only two less SCRs than full-length CFH, and its corresponding cDNA sequences reflects this size difference. Limitations in AAV packaging capacity likely negatively impact expression of the 18tCFH compared to its substantially smaller 12tCFH and FHL-1 counterparts^40^. Therapeutic efficacy of the AAV-mediated tCFH treatment strategies depends on attaining sufficient systemic levels of 18tCFH to prevent uncontrolled C3 activation. Insufficient expression of full-length CFH or any tCFH would not sufficiently dampen uncontrolled C3 activation, enabling subsequent accumulation of the C3 byproducts in the glomeruli. Previous observations of the *CFH-H/0* mouse (C3G+) compared to the *CFH-H/H* mouse (C3G-negative) confirm this pathophysiology, with CFH insufficiency facilitating the development of C3G pathology in the *CFH-H/0* mouse^29^. Without enough CFH or tCFH, C3G and the diseased renal phenotype remains.

Significant differences in the therapeutic efficacy of 12tCFH and FHL-1 underscore the importance of C-terminus SCRs 18-20 in facilitating C3b and GAG binding. Evidence suggests that efficient GAG binding to cell surfaces primarily depends on SCRs 18-20^47^, and this likely contributes to the divergence in 12tCFH and FHL-1’s respective capacities to facilitate C3 clearance in the glomeruli. The absence of critical domains on SCRs 18-20 likely contributes to reduced C3b and GAG binding of FHL-1. Decreases in FHL-1-mediated surface binding and inhibition of C3 activation may also translate to requirements for higher viral titers to achieve therapeutic expression levels. This aligns with previous studies showing that absence of the full-length of the CFH C-terminus prevents surface recognition^48^ functions of the molecule, and therefore likely moderates surface recognition capacity of tCFH as well. Accordingly, requirements for higher viral titers facilitating treatment is shown in **Figure 3** comparing *Cfh-/-* AAV12tCFH to *Cfh-/-*AAVFHL-1 rescue. Analysis of C3G pathology and C3 immunofluorescence in these animals further supports the likelihood of distinct mechanisms underlying efficacy of full-length CFH or tCFH compared to FHL-1. These findings indicate that high doses of FHL-1 are still be capable of rescuing C3G, even if this rescue is less efficient than that mediated by 12tCFH.

Interestingly, the localized therapeutic potential of FHL-1 for rescuing C3G appears to be at odds with previous findings highlighting the inability of FHL-1 to restore ocular complement regulation^25^ following subretinal injection. Initially, structural similarities between the eye and kidney (*i.e.* Bruch’s membrane in the eye and the GBM both contain a network of α3, α4 and α5 type IV collagen chains^49,50^) suggest that both tissues would respond similarly following treatment with AAV8-TBG-FHL-1. However, the treatment of C3G depends on restoration of the APC in the fluid phase, while the treatment of AMD pathology requires a localized response and binding in Bruch’s membrane. For example, significant accumulation of high-density lipoproteins in Bruch’s membrane contribute to AMD^33^, and clearance of these deposits may require surface binding recognition from a therapeutic molecule (*i.e.* 12tCFH, 18tCFH) that outcompetes and ultimately displaces the high-density lipoproteins. In contrast, C3G pathology likely results from uncontrolled C3 activation and its cleavage into C3b, which devolves into excesses of C3b in the fluid phase that are ultimately converted to iC3b^41,51^ before accumulating in the glomeruli; this pathophysiology inherently imposes fewer requirements for surface recognition binding. This interpretation of our findings and of exogenous 12tCFH and FHL-1’s therapeutic capacities direct our consideration of 12tCFH as a top candidate for follow-up studies to develop treatments for C3G. This study provides critical evidence suggesting that AAV-mediated expression of 12tCFH may be sufficient to restore the APC, eliminate C3 deposit accumulation, and resolve the disease burden of C3G.

Our data also supports the high therapeutic potential for the application of an AAV-mediated tCFH Y402 augmentation strategy for the treatment of dry AMD. Strong evidence indicates that the H402 variant of CFH confers significant risk for AMD^15,24,33^, but it remains to be seen whether a tCFH Y402 gene augmentation could successfully rescue an AMD-like phenotype. Future studies to assess this possibility in a mouse model of AMD^33^ will be invaluable our understanding of complement dysregulation in AMD and to the identification of novel treatments. In particular, the capacity of tCFH to clear C3 deposits from glomeruli in C3G suggests that tCFH may be capable of clearing pro-inflammatory accumulates (*i.e.*, high-density lipoproteins^52^, C-reactive protein^53^, and other accumulated molecules) from Bruch’s membrane in AMD. Our data introduces the exciting possibility of AAV-mediated CFH augmentation or replacement as therapeutic strategies, and we establish an important foundation for subsequent efforts to develop novel treatments for patients with AMD and C3G, respectively.

## Materials and Methods

### Generation of Plasmids and AAV Constructs

The 18tCFH cDNA plasmid codes for all 20 SCRs from full-length CFH except SCRs 16 and 17 (**Figure 1A**) and was sequence verified. The 12tCFH cDNA encodes SCRs 1-9 and 18-20 (**Figure 1A**) and was sequence verified. The 18tCFH, 12tCFH, and FHL-1 cDNAs code for the normal Y402 amino acid tyrosine at position 402 (Y402). The thyroxine-binding globin (TBG)– FHL-1–woodchuck hepatitis virus post-transcriptional regulatory element (WPRE) and TBG– 18tCFH plasmids with flanking AAV inverted terminal repeat sequences were generated by VectorBuilder (Chicago, IL, USA) (**Figure 1B**).

The plasmids were packaged into AAV8 capsids as previously described^54^. Briefly, recombinant adeno-associated virus (rAAV) was packaged and purified as previously described^54^. The serotype plasmid AAV8, the transgene containing plasmid flanked by viral inverted terminal repeats (ITRs) [*e.g.*, 18tCFH, FHL-1, and 12tCFH transgenes], and the adenoviral helper plasmid pHelper were transfected in 293T cells (ATCC CRL-3216) using 1mg/ml PEI MAX. 72h after transfection cells were harvested, lysed and the virus was purified using an iodixanol ultracentrifugation gradient. An Amicon Ultra-4 Centrifugal filter unit with a 100kDa MWCO was used for buffer exchange and concentration in sterile PBS. The final viral genome titer was measured by qPCR using primers for the ITRs.

### Animals and Viral Injections

All mice were housed conventionally under ambient light conditions to control for environmental factors and microbiome fluctuations. Mice were maintained in accordance with the Institutional Animal Care and Use Committee at Duke University. The number of mice used for each experiment is provided in the figure legends. *Cfh-/-* mice^26,29^ were generated by Dr. Marina Botto. *CFH-H*, *CFH-H/H*, and *CFH-Y/0* mice were generated, characterized and genotyped as previously described^29^.

For lateral tail vein injections, mice were restrained in a Tailveiner restrainer device (Braintree Scientific, Inc; #TV-150), and tail veins were dilated by contact with a SnuggleSafe heating pad (Lenric C21; #6250). Then, 100 µL of solution containing FHL-1, 18tCFH, or 12tCFH (vg according to each figure) suspended in sterile saline were delivered via a Micro-Fine^TM^ 28-gauge Insulin syringe (#329424; Becton Dickinson).

### Western Blots

Twelve weeks following tail vein injection, mice were euthanized with CO2 and samples collected following transcardial perfusion with phosphate-buffered saline. Samples were prepared and Western blots run as we have previously described^25,33^. Briefly, blood was collected via submandibular vein bleeds in tubes containing EDTA (BD Microtainer; BD Biosciences, Franklin Lakes, NJ) and spun at 4°C to collect plasma. Euthanasia by CO2 and transcardial perfusion followed. Plasma samples were diluted in a 10% 4× stock of XT Sample Buffer (1610791; Bio-Rad, Hercules, CA, USA), with equal volume dilutions corresponding to equal protein quantification. Western blots were run with non-reducing conditions using precast 10% Criterion XT Bis-Tris polyacrylamide gels (3450112; Bio-Rad) and MOPS buffer (1610788; Bio-Rad) with an equal volume of diluted plasma per plasma sample. After transfer, the nitrocellulose membranes were blocked with 10% bovine serum albumin or milk and incubated overnight with primary antibodies [goat anti-CFH (1:10,000–A312; Quidel, San Diego, CA, USA), goat anti-factor B (FB; Kent Laboratories, Bellingham, WA, USA), rabbit anti-C3 (1:5,000 #CSB-PA365507LA01MO, Cusabio). Washes were done with Tris-buffered saline with 0.1% Tween 20 detergent (TBST). Incubation with peroxidase-conjugated bovine anti-goat secondaries (805-035-180; Jackson ImmunoResearch Labs, West Grove, PA, USA) for 1-2 hours was followed by chemiluminescent detection by Pierce ECL (32132; Thermo Fisher Scientific) with a ChemiDoc Imaging System (Bio-Rad). Densitometry was performed using ImageJ software with background subtraction for each lane (National Institutes of Health, Bethesda, MD, USA). Normalization calculations were performed by scaling to expression levels in *CFH-H/H* mice.

### Immunofluorescence and Confocal Microscopy

Mice were euthanized using CO2 and were perfused transcardially with 30 mL phosphate-buffered saline, followed by 4% paraformaldehyde in 0.1 mol/L phosphate buffer (PB), pH 7.4. Kidneys were postfixed in the same fixative overnight and were cut using a vibratome into 50-μm-thick sections. Free-floating sections were placed into glass shell vials and blocked with 0.5mL 10mMEDTA / 0.1% Triton X-100 / 10% normal donkey serum in PBS and incubated for 1 hr at room temperature. The diluent was removed, and the sections were washed with PBS. 0.5mL of goat anti-mouse C3 antiserum (Cat# 55444; MP Biomedicals) at a dilution of 1:500 in PBS was added, and the sections were incubated overnight at 4°C while rocking (Belly Dancer Orbital Shaker, # BDRAA115S; IBI Scientific, Dubuque, IA, USA). The diluent was removed followed by 3 washes in PBS. 0.5mL of the secondary antibody Alexa Fluor TM 488 conjugated donkey anti-goat IgG (H+L) (Cat# A11055, Invitrogen) diluted 1:500 in PBS was then added and incubated for 90 minutes at room temperature while rocking. After multiple washes, DAPI (Cat# 62248, ThermoFisher Scientific) diluted 1:1000 was added to counterstain the nuclei. Images were acquired using a Nikon AXR laser scanning confocal microscope (Nikon Instruments Inc., Tokyo, Japan).

Glomerular immunofluorescence was graded according to whether C3+ immunofluorescent labeling was observed on a given glomerulus. All the glomeruli within an image were counted, and the C3+ fraction of glomeruli for that image was recorded. Overall, 5-15 glomeruli per image (8-12 images per mouse) were analyzed by masked investigators.

### Periodic acid-Schiff (PAS) Staining and Light Microscopy

Kidneys from perfused mice were postfixed overnight in the same fixative before being dehydrated and embedded in paraffin. Ten-micron sections were cut and stained with PAS reagent. Glomerular histologic appearance was graded using the grading system described previously^26^ in which grade 0 was normal; grade I, segmental hypercellularity in 10% to 25% of the glomeruli; grade II, hypercellularity involving >50% of the glomerular tuft in >25% to 50% of the glomeruli; grade III, hypercellularity involving >50% of the glomerular tuft in >50% to 75% of the glomeruli; and grade IV (the most severe), glomerular hypercellularity in >75% or crescents in >25% of the glomeruli. For scoring, 2-3 glomeruli per image (18-24 images per mouse were graded by masked investigators; L.A.C., C.G.H).

## Statistical Analysis

Data were plotted as individual points with error bars representing the standard error (SEM). Unpaired t-tests were used for single comparisons and Tukey’s range test was used for multiple comparisons. *p < 0.05, **p < 0.01

## Acknowledgements

The authors thank Tara Weisz at Duke University for her mouse husbandry work and Amy McArdle at Duke University for instruction ensuring rigorous performance of mouse tail vein injection and submandibular bleeding technique. This work was supported by grants from the National Institutes of Health (NEI R01 EY031748 (CBR), P30 EY005722 (to Duke Eye Center); Foundation Fighting Blindness awards (BR-CMM-0615-0681-DUKE and BR-CMM-0522-0830-DUKE to CBR); and an unrestricted grant from Research to Prevent Blindness (to the Duke Eye Center). Disclosure: L.A. Chew, None; D. Grigsby, None; C.G. Hester, None; J. Amason, None; E.J. Flynn III, None; M. Visel, None; J.G. Flannery, none; C. Bowes Rickman, Applied Genetic Technologies Corporation (F), Aevitas Therapeutics (F), 4DMT (C).

